# Genomic evidence for inbreeding depression and purging of deleterious genetic variation in Indian tigers

**DOI:** 10.1101/2021.05.18.444660

**Authors:** Anubhab Khan, Kaushalkumar Patel, Harsh Shukla, Ashwin Viswanathan, Tom van der Valk, Udayan Borthakur, Parag Nigam, Arun Zachariah, Yadavendradev Jhala, Marty Kardos, Uma Ramakrishnan

## Abstract

Increasing habitat fragmentation leads to wild populations becoming small, isolated, and threatened by inbreeding depression. However, small populations may be able to purge recessive deleterious alleles as they become expressed in homozygotes, thus reducing inbreeding depression and increasing population viability. We used whole genomes sequencing from 57 tigers to estimate individual inbreeding and mutation loads in a small-isolated, and two large-connected populations in India. As expected, the small-isolated population had substantially higher average genomic inbreeding (*F*_ROH_=0.57) than the large-connected (*F*_ROH_=0.35 and *F*_ROH_=0.46) populations. The small-isolated population had the lowest loss-of-function mutation load, likely due to purging of highly deleterious recessive mutations. The large populations had lower missense mutation loads than the small-isolated population, but were not identical, possibly due to different demographic histories. While the number of the loss-of-function alleles in the small-isolated population was lower, these alleles were at high frequencies and homozygosity than in the large populations. Together, our data and analyses provide evidence of (a) high mutation load; (b) purging and (c) the highest predicted inbreeding depression, despite purging, in the small-isolated population. Frequency distributions of damaging and neutral alleles uncover genomic evidence that purifying selection has removed part of the mutation load across Indian tiger populations. These results provide genomic evidence for purifying selection in both small and large populations, but also suggest that the remaining deleterious alleles may have inbreeding associated fitness costs. We suggest that genetic rescue from sources selected based on genome-wide differentiation should offset any possible impacts of inbreeding depression.

**Significance statement:** Habitat fragmentation is sequestering species into small and isolated populations with high chances of extinction. But are small and isolated populations at risk of extinction from inbreeding depression, or does inbreeding and purging of deleterious alleles reduce such threat? Using whole genomes from several wild Indian tiger populations, we provide evidence supporting purging of highly deleterious variants in a small isolated population. However, our analyses also indicate that the remaining highly deleterious alleles are at high frequencies, suggesting continued inbreeding depression despite some successful purging. We discuss the implications of our results for conservation, including possible genomics-informed genetic rescue strategies.

## Introduction

A large proportion of the earth’s biodiversity persists in small and isolated populations in today’s anthropogenically modified world^1^. Such populations may suffer from decreased genetic variation and increased inbreeding^2^ which together lead to decreased fitness and increased extinction risk^3^. Several theoretical^4,5,6,7^ experimental^8^ and empirical studies (for example 9) reveal that species surviving in small and isolated populations are at the greatest risk of extinction.

While species exist in nature along a continuum from small to large populations with different levels of isolation, populations of endangered species often tend to be small and isolated. African wild dog, Ethiopian wolf, and great Indian bustard are examples of species where all populations are small and isolated^10,11,12^. The ‘small population paradigm’ of conservation biology suggests that such smaller and more isolated populations are at a higher risk of extinction due to inbreeding depression and demographic stochasticity^13,14,15^.

Inbred individuals express deleterious, partially recessive alleles that are inherited identical-by-descent (IBD) from related parents, leading to inbreeding depression^16^. Such inbreeding depression can reduce the average fitness of a population, eventually leading to reduced population size, and possibly extinction^17^. A commonly adopted strategy to conserve inbred populations is genetic rescue^18^, which aims to increase average fitness by decreasing the frequency of deleterious mutations and increase heterozygosity at loci harboring deleterious alleles, via translocations of individuals from genetically differentiated populations. A meta-analysis of empirical data from wild populations showed broadly consistent positive effects of genetic rescue on fitness^15,19^.

Population genetic theory^20,21,22^ predicts that purifying selection can reduce inbreeding depression by purging deleterious alleles from inbred populations in the absence of immigration. Whether isolated populations are likely to purge a substantial fraction of the mutation load has been of longstanding interest in evolutionary biology and conservation. Early empirical data from pedigreed captive populations suggested that purging was either absent, or resulted only in slight decreases in inbreeding depression (e.g., 23,24). However, several experimental studies based on model organisms reveal substantial purging and significant reduction in inbreeding depression in small populations25,26,27. Recent molecular and population genetic studies have found genomic evidence for purging in wild populations (e.g., 28,29,30). Despite broad empirical support for the efficacy of genetic rescue^15,19^, genomic evidence for purging and the long term persistence of some small isolated populations have been cited to question the small population paradigm, and to argue that standard genetic rescue practices are likely to be counterproductive^29,31,32,33^. Whether purging removes enough deleterious alleles to improve the viability of small, isolated populations (contradicting the small population paradigm) remains an open question. We address this question by contrasting genomic inbreeding and mutation load in small isolated versus large-connected populations of wild tigers.

We use Bengal tigers (*Panthera tigris tigris*) from India as a model to investigate levels of inbreeding and relative mutation loads in small-isolated and large-connected populations, and examine the potential for genetic rescue. Tigers are large, endangered carnivores, but Bengal tigers have high genetic variation compared to other sub-species, with some sub-populations showing high inbreeding indicating isolation^34^. All Bengal tiger populations have been through historic bottlenecks, but inbreeding and genetic variation vary among populations. Also, some populations are relatively large and connected while others are small and isolated from both genetic^35^ and demographic perspectives^36,37^, making them an ideal system to investigate inbreeding, mutation load and possible genetic rescue strategies.

We use genomic data to measure the impact of historically declining population sizes and connectivity on inbreeding and mutation load in three wild Bengal tiger (*Panthera tigris tigris*), populations that are small-isolated (SI) and large-connected. The small-isolated population is from north-western India and the large-connected populations are from south India (s-LC) and central India (c-LC). We expected large-connected populations to be the least inbred and to have the lowest mutation load. Alternatively, purging could result in lower mutation load in this small-isolated population. We also explore strategies for genetic rescue that might effectively decrease inbreeding depression. For tigers, we specifically suggest strategies for identifying populations that may benefit from genetic rescue, and how such strategies may be effectively implemented.

## Results and Discussion

### Genomic estimates of inbreeding

Tigers exist as several genetic population clusters within India with the total population size of these clusters ranging from 60 to 1000 individuals. Each genetic cluster consists of several protected areas, with varying connectivity with other such protected areas and clusters^35,36,38^. For the purpose of this paper, we defined populations, and whether they are ‘isolated’ or ‘connected’ based on population genetic and gene flow analyses using markers across the genome^34,35,39^. Here, the ‘large’ populations currently have hundreds of tigers (minimum 300 tigers), while ‘small’ populations currently have below 100 tigers^36^. While both large populations are part of connected landscapes, they do not have identical histories of connectivity. The southern large-connected population is disconnected from other tiger population genetic clusters in India (e.g. from Central India and Northwest), while central large-connected population (Central Indian landscape) was connected by gene flow to other tiger genetic clusters until recently (see 39,35). SFS-based demographic history models in Armstrong et al.^34^ suggest that most Indian tigers diverged from each other recently. Increased agriculture, bounty hunting and illegal poaching, habitat loss and fragmentation together have led to local extinctions of tiger populations^40,41^. The small-isolated population we studied here has less than one hundred individuals, but was connected (until very recently) to the central large-connected population^42^ and potentially to the now extinct tiger population in Afghanistan^43^.

We quantify inbreeding by identifying long stretches of the genome that are homozygous and identical by descent (i.e., runs-of-homozygosity, ROH). The summed length of ROH divided by total autosomal size is an accurate genomic measure of inbreeding (*F*_ROH_) (e.g., 44). We found that the mean *F*_ROH_, measured using ROH longer than 100Kb (*F*_ROH>100Kb_), 1Mb (*F*_ROH>1Mb_) and 5Mb (*F*_ROH>5Mb_) of all individuals in our dataset (regardless of population) was above 0.48, 0.31 and 0.23 respectively. Tigers from the small-isolated population were most inbred with an average *F*_ROH>100Kb_ of 0.57, while average inbreeding due to recent ancestors (on average up to 5 generations ago, contingent on assumed recombination rate from domestic cat^45^) was above *F*_ROH>5Mb_= 0.37 (Figure 1a). The *F*_ROH_ was proportional to effective population size (*N*_e_), with the small isolated population having lower *N*_e_ at all time intervals compared to both large and connected populations. This level of inbreeding is empirically associated with high risk of extinction^46,47,48^. However, shorter ROH could be due to shared demographic history of bottlenecks (e.g. *F*_ROH_=100kb) and inbreeding depression may be observed at higher *F* values for large carnivores^49^. Pedigree based studies^50^ have reported that an inbreeding coefficient of 0.47 is lethal for tigers.

**Figure 1:**
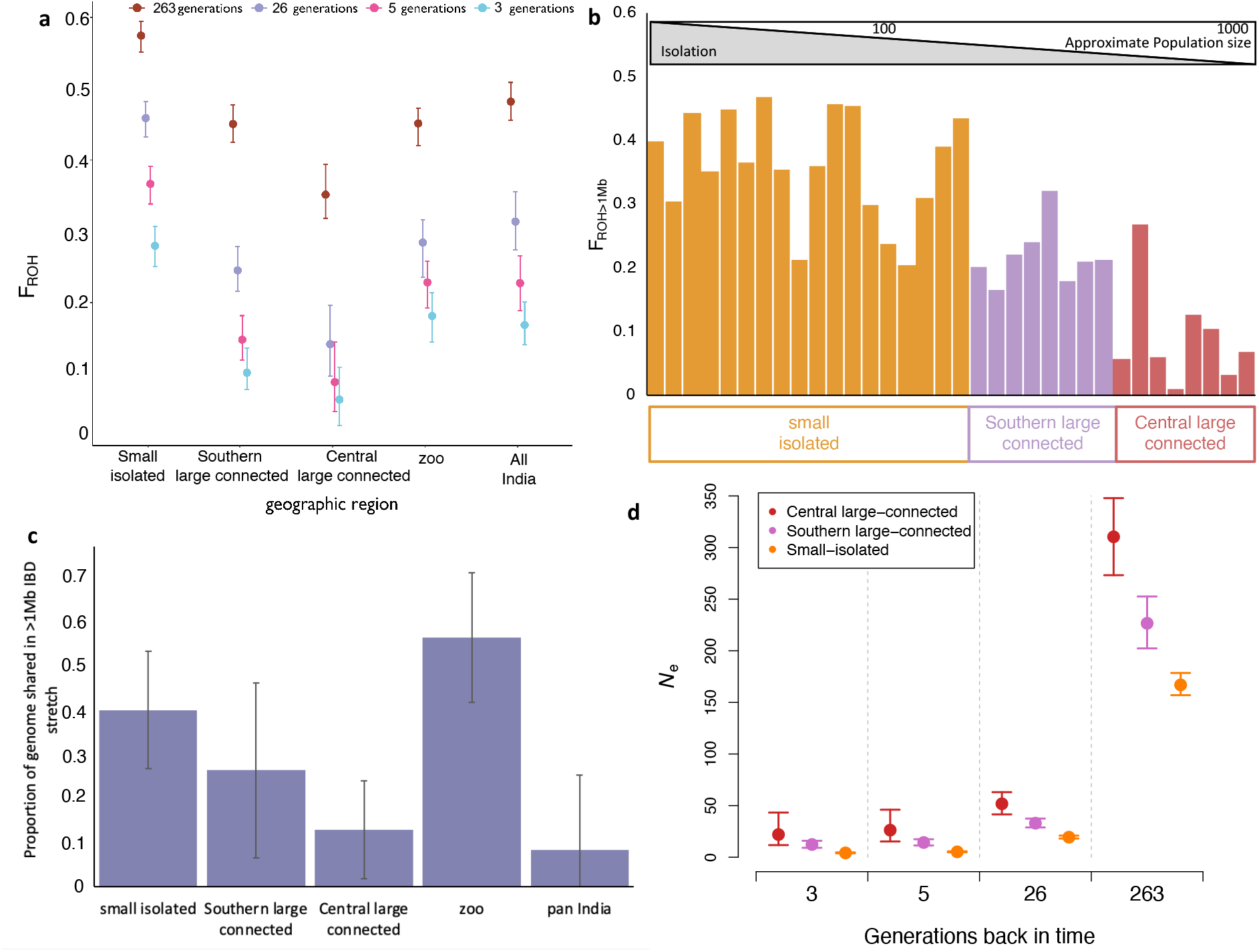
Inbreeding and shared ancestry in Indian tigers. Here we show cumulative inbreeding due to parental shared ancestry due to ancestors at different historical time ranges as estimated by different lengths of ROH, that is, ROH ≥ 10 Mb indicate shared ancestry on average up to 3 generations ago, 5 Mb stretches indicate on average up to 5 generations, 1 Mb stretches indicate on average up to 26 generations and 0.1Mb stretches indicate on average up to 263 generations ago. The figure shows cumulative inbreeding in each population due to shared ancestry between parents at each time range. Error bars are 95% CI. (a), inbreeding in individuals due to shared ancestry 26 generation ago (b), the proportion of the genome shared in in 1 Mb or longer tracts between pairs of individual from a population of individuals belonging to central large connected, southern large connected and small isolated populations. Pan India comparisons in (c) include all pairwise comparisons of wild individuals. Error bars are standard errors. *N*_e_ over time as estimated from *F*_ROH_ for the different populations. Error bars are standard deviations (d).

We show that inbreeding was low in the large-connected populations compared to the small-isolated population (Figure 1a,b). We compared our estimates for zoo individuals with their known pedigree values to authenticate these results and found that estimates of inbreeding indeed correspond with pedigree values^51^ (Supplementary Table 1). Overall, the long ROH of ≥10Mb, signifying inbreeding due to ancestors as recent as 3 generations, are rare in large populations but frequent in the small-isolated population. The mean inbreeding due to recent ancestors in the small-isolated population (*F*_ROH>10Mb_) was above 0.28, which could lead to several negative effects^46,48^. We found that the large connected populations showed inbreeding due to recent ancestors (on average up to 5 generations ago) of less than *F*_ROH>5Mb_ = 0.1. The large southern population has more inbred individuals than the large central population, potentially due to its geographical placement, historic size (Figure 1d) and connectivity^35,36,39^. Overall, although the theoretical expectation is that populations with *N*_e_ < 1,000 are unlikely to persist^7,30,52,53^, this result indicates that a population with a census size of ∼600 tigers may be sufficiently large to buffer short term inbreeding effects with some geneflow. The long term viability of these populations remains to be tested. While there are instances of small tiger (and other carnivore) populations persisting^36,37^, this could possibly be due to their ability to purge some deleterious alleles. This implies that other large felid species, like lions^54^, snow leopards^55,56^ and jaguars^57^ that often persist in population sizes close to 600 may not necessarily experience the detrimental effects of inbreeding depression over the short term. We caution that this is likely to depend on founder effects and chance variation in the mutation load among populations (but see 30), and does not address the likelihood for fitness decline over the long term^7^.

In addition to having the highest *F*_ROH_, we found that pairs of individuals from the small-isolated population also shared large tracts of genome that were IBD (Figure 1c). This could be a worrying sign for its long-term future as the pairwise IBD sharing translates into inbreeding in the offspring. On average, pairs of individuals from the small-isolated population shared about 40% of their genome (as more than 1Mb long IBD stretches), while pairs from the southern large-connected and the central large-connected populations share about 25% and 15% respectively, slightly higher than the corresponding mean *F*_ROH_ values (Figure 1a). As with inbreeding in offspring, pairwise IBD sharing is expected to be influenced by founding bottlenecks in addition to recent small *N*_e_ (Figure 1d). The small-isolated population had the lowest mitochondrial haplotype diversity and the lowest number of mitochondrial haplotypes (Supplementary Figure 1). This lack of diverse lineages, large tracts of shared IBD genome between individuals, and the recent small population size may therefore require continued immigration of individuals to sustain genetic variation^58^, or in the absence of natural immigration, may require genetic rescue.

### Mutation loads in small-isolated and large-connected tiger populations

We evaluated the relationships between *F*_ROH_ and the frequency and genomic distribution of putatively deleterious genotypes identified using the Ensembl Variant Effect Predictor^59^ (VEP). Under the assumption that most deleterious alleles are partially recessive^28,60,61^, the number of homozygous damaging alleles is likely to be informative of the fitness cost of individual inbreeding. First, we found that the number of homozygous putatively damaging alleles (loss-of-function (LOF) and missense alleles together) was proportional to *F*_ROH_ in all populations. The number of putatively damaging homozygotes increased from <1500 for individuals with *F*_ROH_ ≈ 0.3 to approximately 2200 for individuals with *F*_ROH_ ≈ 0.6 (*P* < 0.01 for each population, linear regressions; Figure 2). This genomic evidence suggests a fitness cost associated with increased inbreeding. Additionally, the burden of homozygous putatively damaging mutations was substantially larger on average in the small-isolated population (mean ≈ 2,000 per individual) than in the central large-connected population (mean < 1,600 per individual, *P* < 0.0001 randomization test) or the southern large-connected population (mean > 1,800 per individual, *P* = 0.0025, randomization test, Figure 2). This is likely due to higher *N*_e_ in the central large-connected population as compared to the southern large-connected population (Figure 1d).

**Figure 2.**
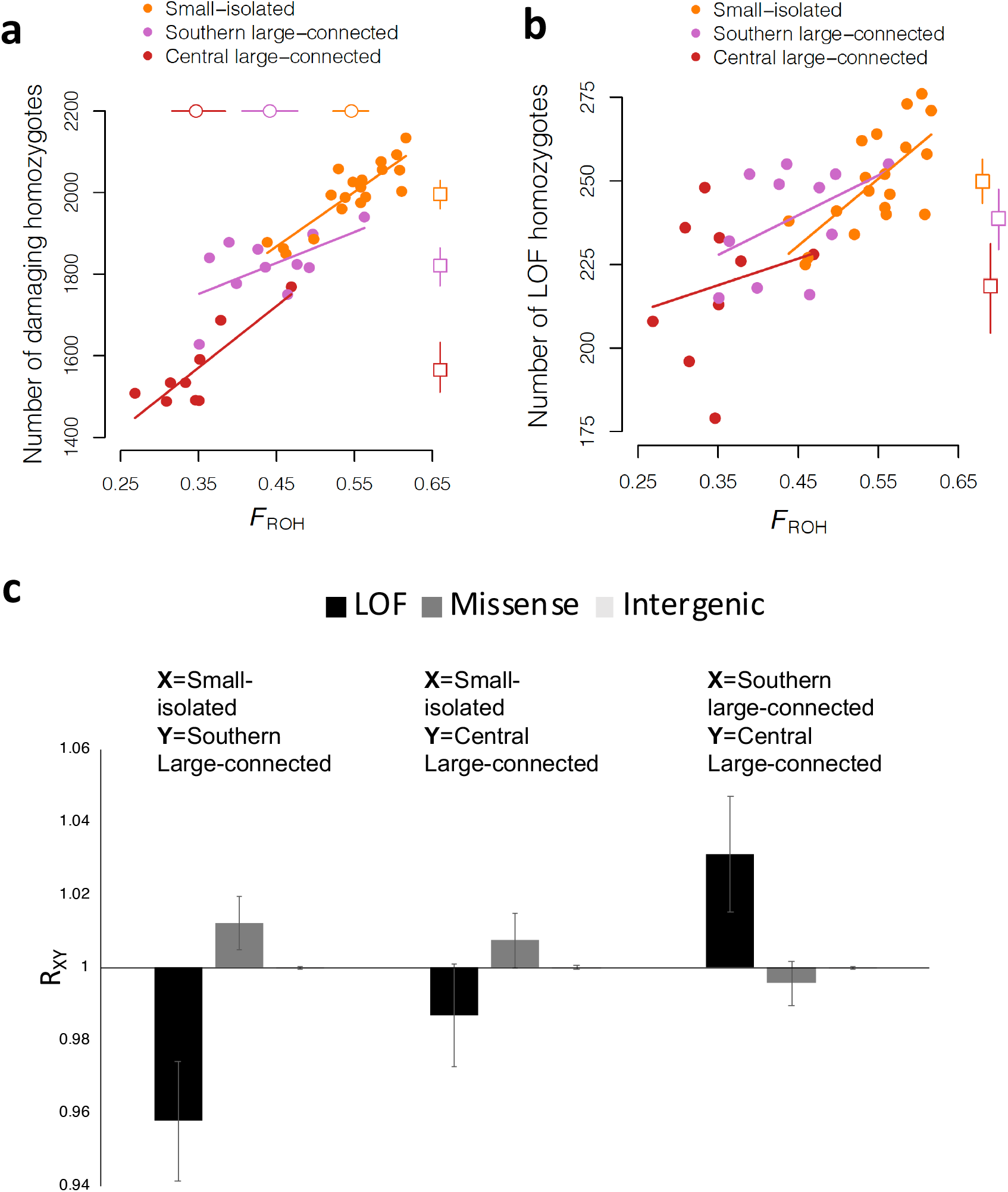
Relationships between inbreeding and number of homozygous damaging alleles (LOF plus missense) **(a)**, and homozygous LOF mutations **(b)**, and relative mutation loads in populations of various sizes and levels of isolation **(c)**. Filled circles represent the values of *F*_ROH_ (x-axis) for different individuals, and the number of homozygous damaging mutations (y-axis), with each population represented by color as indicated in the legend. Open circles and squares represent the mean *F*_ROH_ and mean number of homozygous damaging mutations, respectively, for each population, with the error bars representing 95% percentile bootstrap confidence intervals. R_XY_ indicates excess of deleterious alleles in population X relative to Y. All populations had similar numbers of neutral (intergenic) mutations, and error bars are standard errors from jacknife in **(c)**.

We further investigated differential mutation load between populations for LOF and missense mutations identified with VEP, and the R_XY_ method^28,62^ to estimate relative excess of number of putatively deleterious alleles in one population compared to another. R_XY_ compares the number of derived alleles of a particular category (LOF, missense, or putatively neutral) found in one population relative to the other. If R_XY_ is one, both populations X and Y have the same load of derived alleles, whereas if R_XY_ > 1 then population X has more derived alleles than Y and vice versa if R_XY_ < 1 ^28,62^. As above, we found a higher load associated with missense mutations in the small-isolated population (Figure 2c). This is likely explained by more efficient purifying selection against missense mutations of relatively small effect, which are typically only slightly recessive^61^, due to larger *N*_e_ in the large populations than in the small population. We also found a lower load of LOF mutations in the small-isolated population compared to the large populations (Figure 2c). This suggests that higher inbreeding in the small-isolated population facilitated purging of relatively large-effect LOF mutations – which are more likely to be highly recessive than small-effect deleterious mutations^61,63^. Together, these different approaches suggest a higher putatively damaging allele load in small isolated population coupled with purging of loss-of-function alleles in the small isolated (and inbred) tiger population.

Despite this evidence for purging of LOF alleles in the small-isolated population, it appears that high inbreeding lead to individuals in this population having a higher number of homozygous LOF alleles on average (N=250) compared to the southern large-connected (N=238) and central large-connected (N=218) populations (Figure 2b). High homozygosity for remaining LOF alleles suggests that the fitness cost due to LOF mutations may be higher in the small-isolated population compared to the large populations despite partial purging. Importantly, while it is impossible to know the realized fitness effects of homozygosity for LOF and missense mutations, this analysis suggests that purging was not sufficient to result in a smaller fitness cost of inbreeding in the small-isolated population compared to the larger populations. We observe similar results when we used a Genomic Evolutionary Rate Profiling (GERP)^65^ analysis to identify potentially highly deleterious alleles by keeping loci with GERP scores in the top 0.1 percent (Supplementary Figure 3). For this analysis, and those below, it important to recognize that our measure of mutation load is not a direct measure of fitness. Rather, our inferences rest on the assumption that missense and LOF mutations, and mutations at conserved sites are deleterious on average, which is well supported empirically^65,66^. Definitively determining whether the bottlenecks faced by these populations^40,41^ and the additional bottleneck faced by north-west Indian population three generations ago in 2004^67^ purged part of the mutation load would require historical genetic data to quantify pre-versus post-bottleneck mutation loads. Additionally, better estimates of very recent demographic history would help to predict the potential for purging during the bottlenecks.

Annotated sets of deleterious variants are rarely available for endangered species. We compared our missense mutations to annotated sets of domestic cat mutations (Supplementary Table 2). The missense mutations we identified mapped to genes responsible for disease states such as hypertrophic cardiomyopathy, progressive retinal atrophy, polycystic kidney disease, cystinuria, gangliosidosis, hyperoxaluria, hypothyroidism, mucolipidosis and mucopolysaccharidosis. Further studies are needed to functionally validate the effect of these mutations in tigers. Studies on inbred/small populations of Asiatic lions revealed several abnormalities^68^, and genomic studies here might provide further insights on deleterious mutations in large cats. In the small-isolated population at least one tiger has a potential eye condition (Supplementary Figure 2), although no skeletal defects were observed in an earlier study^69^. Additional quantitative genetic studies on inbred zoo individuals can aid in understanding the possible phenotypic effects of inbreeding and inbreeding depression for endangered species.

Site-frequency spectra (SFS) revealed that nearly 14% of both putatively damaging and neutral derived alleles were fixed in the small-isolated population (Figure 3). A smaller proportion of derived alleles (both damaging and neutral) were fixed in the large populations (< 0.06). However, there were slightly more fixed derived alleles in the southern large connected population than in the central large-connected population. The higher abundance of fixed derived alleles (both damaging and neutral) in the small-isolated population, and to a lesser extent in the southern large-connected population, suggests that genetic drift (e.g., due to historical population bottlenecks and founder effects) drove previously rare alleles to fixation in these populations (Figure 3)^70^. The fixation of deleterious alleles is consistent with theoretical predictions for very small populations^4,7^ and empirical observations from wolves^32^, Apennine brown bears^71^ and other species^31^. In the Isle Royale wolves, high frequency deleterious alleles were consistent with frequent bone deformities in the population^32,72^. The large fraction of fixed putatively damaging mutations in the small isolated population suggests that partial purging of the mutation load due to LOF mutations (Figure 2c) was not sufficient to erase a substantial fraction of the mutation load in this population. While the SFS was essentially flat for polymorphic loci (i.e., excluding loci with derived allele frequencies of 0 or 1) in the small isolated population (as expected following a population bottleneck), there was a larger fraction of rare derived alleles in the large populations, as expected for large populations at mutation drift equilibrium^70^. A larger proportion of putatively damaging than neutral alleles were at or near fixation in each of the large populations, and across all of India (Figure 3), possibly due to positive selection driving selective sweeps on a small fraction of non-synonymous mutations. The hump-shaped SFS are likely the result of a complex demographic history for Indian tigers, including strong population structure, and historical population bottlenecks^70,73,74^.

**Figure 3.**
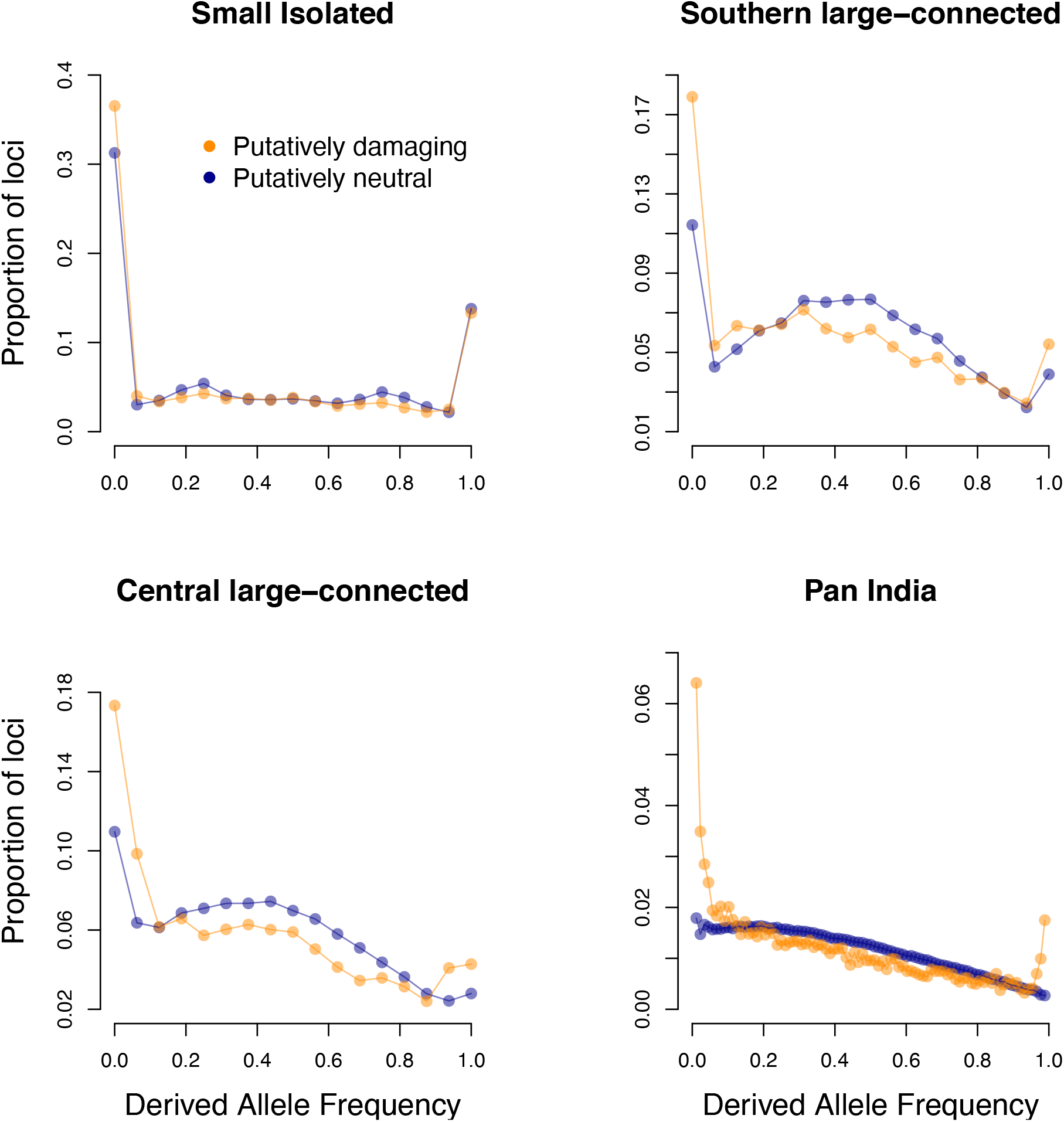
Site frequency spectrum for damaging (missense and LOF mutations combined, orange) and neutral mutations (blue) for each population. The proportion of loci (y-axis) is shown for each possible derived allele frequency (x-axis). Sample sizes were equalized among the three populations by randomly subsampling 16 non-missing alleles for each locus in each of the three populations. Fixed (frequency=1) and missing (frequency=0) alleles are included in the SFS for the three individual populations. We subsampled 47 non-missing alleles at each locus to calculate the SFS for pan India to account for missing genotypes.

Lastly, the frequency distributions of putatively deleterious and neutral derived alleles reveals signatures of purging via natural selection. Some previously rare, deleterious alleles are expected to be removed from a population by random genetic drift alone, during periods of small *N*_e_ (e.g., population bottlenecks or founding events). Effective natural selection through purging is expected to eliminate deleterious derived alleles more frequently than neutral alleles, or to keep them at lower frequencies than neutral derived alleles. We expected putatively deleterious derived alleles to either be absent, or to be less frequent (on average) than putatively neutral derived alleles if purging has occurred in our study populations. This prediction was supported via a randomization test in each population and for all populations combined (*P* < 2e-4, 2-sided randomization test). This suggests that purifying selection has successfully removed a small fraction of the mutation load in each of the three study populations and across India. The higher proportion of fixed and very high frequency derived putatively damaging alleles in the large populations and across Pan India may be caused by historical selective sweeps on a small fraction of positively selected nonsynonymous mutations, the details of which we leave to future research.

Interestingly, both inbreeding and mutation load for the southern large connected population (from Western ghats in India) are higher than the central large connected population (from Central India, Figures 1 and 2). This could reflect the central location of this population (see 35,38,39). Additionally, the central large connected population may have had higher historical connectivity (with other population clusters), a two dimensional network of local populations (versus a linear array in southern large connected populations) and other ecological factors that allow higher population densities to be achieved in all habitats of central large connected population compared to southern large connected populations.

### Genetic rescue strategies

Our data suggest that genomic inbreeding (*F*_ROH_ up to 0.6 in the small isolated population) is comparable to populations experiencing inbreeding depression (e.g. Florida puma, *F*_ROH_ up to 0.6^75^ and Isle Royale wolves, *F*_ROH_ between 0.1 and 0.5^32^). While we also demonstrate the effects of purifying selection, homozygous LOF alleles are more frequent, and mildly deleterious missense mutations are in excess in the small isolated population.

Gene flow is the proposed strategy to sustain small and isolated populations^18^. Geneflow that results in masking of deleterious alleles responsible for genetic load, leading to increased population growth rate is genetic rescue. Evolutionary rescue is geneflow that increases genetic variation^76^ and thus adaptive potential. High *F*_ST_ at loci with high frequency damaging alleles in the receiving population (frequency > 0.9) should be indicative of good source populations for genetic rescue. In practice, for both evolutionary rescue and genetic rescue, source populations are identified as those that have high genome-wide genetic differentiation with the receiving population (to maximize the heterozygosity of admixed offspring (reviewed in 19)). Geneflow from populations with high genome-wide *F*_ST_ in theory should result in masking deleterious alleles, allowing both genetic rescue and evolutionary rescue. However, the smaller number of loci with high-frequency damaging alleles compared to all polymorphic loci could lead to discordance between these two proxies for genetic and evolutionary rescue (i.e., genetic differentiation at loci with putatively damaging alleles may not be a good predictor of genome-wide genetic differentiation).

We investigated whether evolutionary rescue is a good proxy for genetic rescue. We propose that certain source populations may be ideal if they result in masking of damaging alleles and increased genome-wide heterozygosity (top right quadrant of Figure 4a). Alternatively, other source populations (bottom right quadrant) will increase heterozygosity but not mask damaging alleles, while those in the top left quadrant mask damaging alleles but may not increase heterozygosity substantially genome-wide (Figure 4a). Our data and estimates reveal a strong and positive correlation between the two proxies (Figure 4b, adjusted *r*^2^ = 0.91), re-enforcing that selecting source populations based on high genome wide *F*_ST_ would result in both genetic and evolutionary rescue. For the small isolated population, our data suggest that Kaziranga (proportion of damaging loci rescued = 0.74; Genome-wide *F*_ST_ = 0.26, damaging loci *F*_ST_ =0.26) is the best source population, while ecologically more similar populations like Kanha (proportion of damaging loci rescued = 0.69; Genome-wide F_st_ = 0.22; damaging loci *F*_ST_ =0.23) and Corbett (proportion of damaging loci rescued = 0.65; Genome-wide F_st_ = 0.26; damaging loci *F*_ST_ =0.28) may be more practical, and almost as effective at genetic rescue. For the large isolated population in Southern India, both proxies are lower on an average (average proportion of damaging loci rescued = 0.65; average genome-wide *F*_ST_ = 0.195, average damaging allele *F*_ST_ = 0.21, across source populations). Note that high genome-wide *F*_*ST*_ between a donor and recipient population could also be due to intense bottlenecks and resulting drift in endangered species, and such population pairs may not result in effective genetic rescue. We caution that these inferences are subject to assumed fitness effects of damaging alleles. Most genome-wide predictions of deleterious alleles in wild species await annotation with phenotypic data^77^, such a strategy could be implemented when the loci affecting fitness are identified as in the case of Tasmaninan devils (*Sarcophilus harrisii*^78^) and Soay sheep^79^.

**Figure 4.**
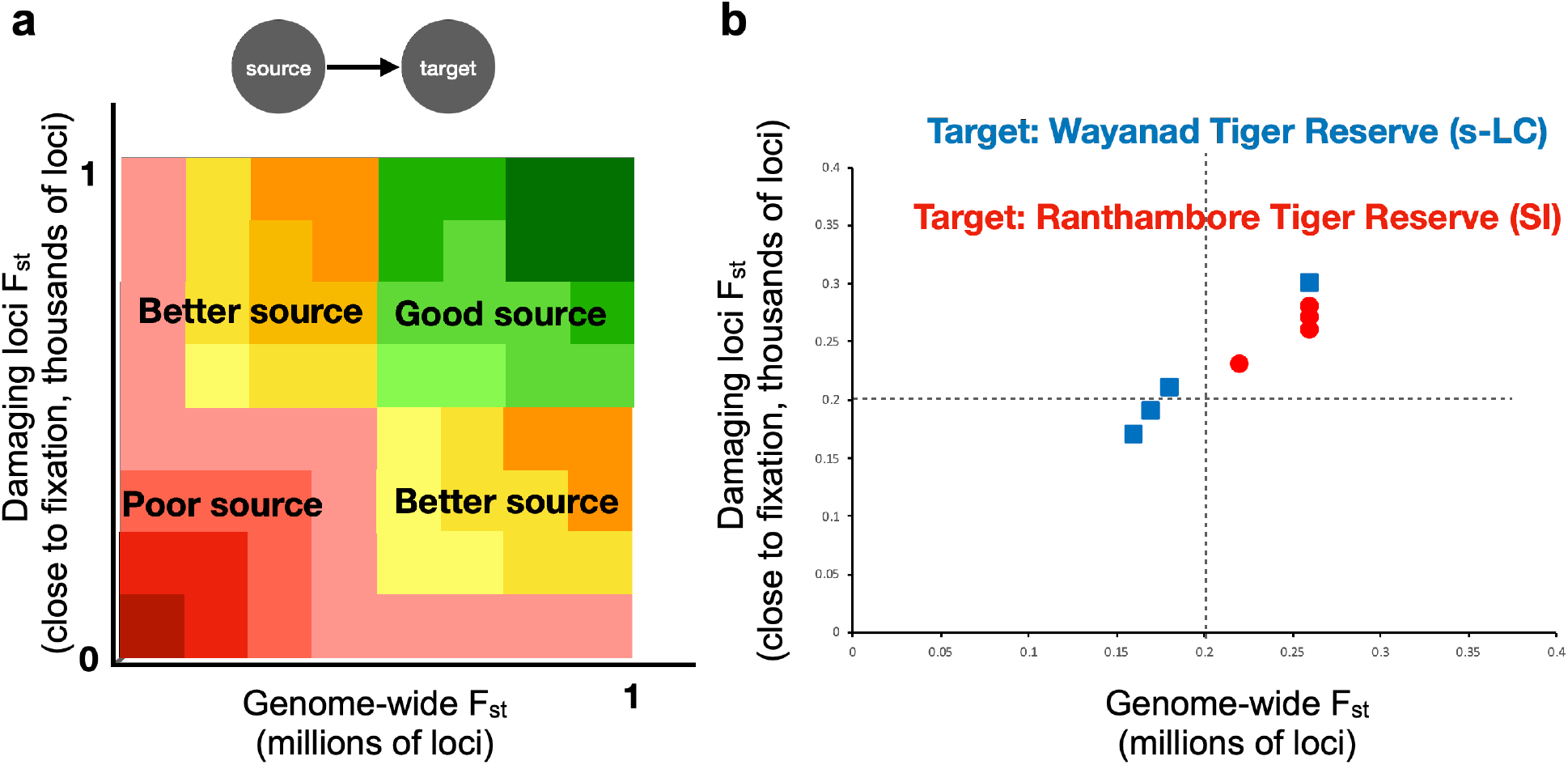
(a) A schematic showing the possible distribution of genome-wide *F*_ST_ and (b) the observed values for our data, assuming two target populations, Ranthambore Tiger Reserve (in red circles, the small isolated population) and Wayanad Tiger Reserve (blue squares, a protected area within the Southern large connected population). *F*_ST_ on the y-axis is for loci where the putatively damaging allele is at a frequency >0.9 in the target population, and for all loci across the genome on the x-axis.

Whether there is ever a substantial advantage to selecting source populations for genetic rescue attempts based on differentiation for putatively damaging loci (versus across the whole genome) remains to be tested empirically. Genome wide surveys would help in identifying source populations that best mask damaging alleles and increase genome-wide heterozygosity; however there are other considerations (as highlighted in 33) that could impact selection of source populations. A denser sampling of tigers across the Indian tiger range is necessary to predict best rescue individuals for particular target populations. Such dense sampling of genomes can benefit from sequencing of non-invasive samples since acquiring blood samples from endangered species can be difficult^80^. Methods involving sequencing genomes from shed hair^81^ or fecal samples^82^ will be useful in creating such translocation networks and monitoring the populations for gene flow.

### Implications for conservation of endangered species

We find that small and isolated populations, in this case tigers of Ranthambore Tiger Reserve, have high inbreeding arising from both recent ancestors and ancestors in deep history. Our analyses revealed a signature of purging for putatively large effect (LOF) deleterious alleles in this population. However, this small and isolated population also had the highest average number of homozygous damaging alleles of the three study populations, suggesting potential fitness cost of inbreeding in this population despite purging of some large effect mutations, and compared to the large and connected populations. Together, these results demonstrate that purging (as inferred from genomic signatures) does not eliminate all damaging alleles, and hence does not contradict the small population paradigm of conservation biology^19^. Careful predictions of putative phenotypic effects of the existing deleterious alleles at high frequencies might be instructive to understand future inbreeding depression, and these alleles could potentially be used as early warning signs for population decline. However, caution is warranted in this regard because it is still unclear how effectively molecular predictions (i.e., identification of loss of function or derived alleles at conserved sites) can translate into predictions of fitness differences among extant individuals and populations.

Historical isolation and bottlenecks due to overharvest appear to have a significant effect on missense (potentially mild effect) mutation load. Our analyses suggest that purging can also occur in large-connected populations in the context of endangered species (Figure 3). This could be because all tiger populations are in general small, and the range of population sizes is not very wide^36^. Continued isolation of these populations will increase mutation load in the future. Thus maximizing connectivity might be the best strategy to minimize extinction (from an ecological perspective), increase genome-wide genetic variation, and also minimize mutation load in the future. Maximizing connectivity in the wild would require increasing population abundance since large populations are sources of dispersing individuals, but also planning for habitat corridors, since demographics of tiger landscapes are dominated by extinction and recolonization dynamics (e.g. see 83, 84). Such strategies may be relevant to several endangered species with wide ranges but fragmented populations, like elephants, lions and wild dogs.

Critical to providing recommendations for management is understanding ongoing evolutionary trajectories for endangered species. Given the large number of frequent deleterious alleles in the small isolated population, management could aim to decrease the number of close to fixation damaging alleles, potentially maximizing mean absolute fitness. Our data reveal that source populations with high genome-wide differentiation (with the target population) would adequately mask deleterious alleles and increase heterozygosity. Gene flow from even the relatively proximate large-connected population in Central India would decrease the frequency of damaging alleles, and allow genetic rescue. We caution that such gene flow could also introduce other damaging alleles (e.g. in wolves, see 85).

Overall, genomic data and analyses provide richer, more nuanced ways to address inbreeding and genetic rescue of small and isolated populations of endangered species (i.e., the small population paradigm in conservation). We demonstrate effective purging of some loss-of-function alleles, but not mutations with smaller effects, highlighting the need to characterize detrimental mutations and how they will be addressed through genetic rescue in future conservation efforts. Conservation management based on predictive models for fitness effects in the context of observed mutations will aid long-term persistence of populations in the wild.

## Methods

### Sample collection

We collected tissue from tranquilized or dead tigers from several tiger reserves (14 protected areas across India); zoo individuals from Sagar et al.^51^ were also analyzed (Figure 5, Supplementary Table 1). Individuals belonged to one of five population clusters: 1) **Small-isolated population** from north-western India (NW) (n=19) **-** we obtained tissues from 15 NW individuals and used data from two NW individuals in Armstrong et al.^34^ and two NW individuals in Khan et al.^81^. Our NW samples consist only of individuals that were born in Ranthambore Tiger Reserve (RTR) but could have been translocated to other protected areas (for example Sariska Tiger Reserve) as per forest department records. 2) **Central Large-connected population** from central India (C) (n=9) **-** we obtained tissue from four Kanha Tiger Reserve (KTR) individuals and used data from Armstrong et al.^34^ for five other KTR individuals. 3) **Southern Large-connected population** from southern India (n=11) **-** we obtained tissue from two Wayanad Wildlife Sanctuary (WWS) individuals and two Bandipur Tiger Reserve (BTR) individuals. We additionally used data from Armstrong et al.^34^ for five WWS individuals and Liu et. al.^86^ for two Nagarhole Tiger Reserve (NTR) individuals. Since WWS, BTR and NTR populations are effectively contiguous, we refer to them together as Wayanad (WAY) tigers. 4) **Nandankanan Zoo**, Odisha, India – Sagar et al.^51^ obtained five blood samples from inbred individuals from Nandankanan Zoo, and these genomes were used as a control to test our methods. The pedigree based inbreeding coefficient of these individuals vary from 0.21 (n=1), 0.26 (n=1) and 0.28 (n=3) (Supplementary Table 1). Such pedigree-based estimates are expected to be an underestimate of the actual inbreeding due to the low depth of pedigrees^44^. 5) **Other protected areas -** we also sampled one individual tiger from Lalgarh Division Forest, two individuals from Corbett Tiger Reserve (COR), one from Chitwan National Park, four from Kaziranga Tiger Reserve, one from Periyar Tiger Reserve, one from Bor Tiger Reserve, one from Chandrapur district in Maharashtra and two from Sundarban Tiger Reserve. These individuals combined with others are used for drawing pan-India inferences. The sampling locations are depicted in Figure 6.

**Figure 5.**
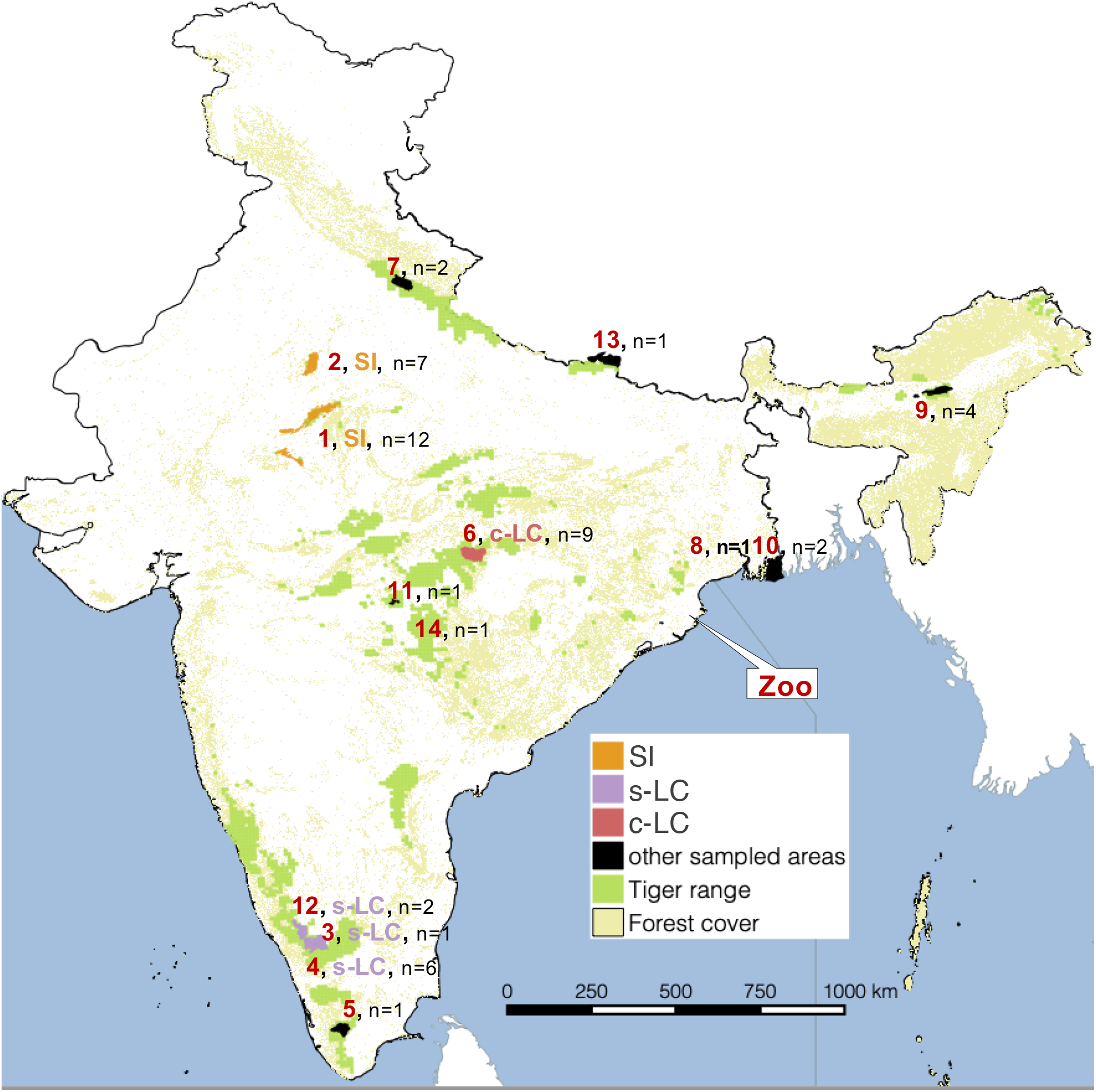
Sample locations and populations. The numbers in the figures represent the protected area where, North-west population: 1=Ranthambore Tiger Reserve (n=11), 2=Sariska Tiger Reserve (n=7), South India population: 3=Bandipur Tiger Reserve (n=3), 4=Wayanad Wildlife Sanctuary (n=6), 12=Nagarhole Tiger Reserve (n=2), 5=Periyar Tiger Reserve (n=1), Central Cluster: 6=Kanha Tiger Reserve (n=9), 7=Corbett Tiger Reserve (n=2), 8=Lalgarh Range (n=1), 10=Sunderban Tiger Reserve (n=2), 11=Bor Tiger Reserve (n=1), 13= Chitwan National Park (n=1), 14=Chandrapur (n=1), North East population: 9=Kaziranga Tiger Reserve (n=4), Zoo: Zoo=Nanadankanan Zoo (n=5), Sagar et al. 2021.

### DNA extraction and sequencing

We extracted DNA using Qiagen blood and tissue extraction kit (Cat. 69504) as per the manufacturer’s instructions. We prepared DNA whole genome sequencing libraries using NEBNext UltraTM II DNA Library Prep Kit (Cat. E7645L, NEB Inc.). We quantified DNA on QubitTM 3.0 fluorometer using Qubit High sensitivity dsDNA Assay (#Q32854, Thermo Fisher Scientific). We then fragmented the quantified DNA by sonication using Covaris LE220 ultrasonicator and Covaris microTUBE (#520053, Covaris® Inc.) to obtain a final insert size of 250–350 bp. Next, we performed end-repair on the DNA fragments where blunt ends are created on either side of the fragments. We added a single “A” nucleotide on the 3′ ends of the fragments to facilitate the ligation of NEB stem-loop adapters. We cleaned the ligated products and size selected using Agencourt AMPure XP beads (#A63882, Beckman Coulter). We amplified these size selected products using an eight-cycle PCR during which indices (Barcodes) and flow-cell binding sequences were added. After a final clean-up with Agencourt AMPure XP beads, we quantified the libraries using Qubit DNA Assay and assessed the fragments using DNA TapeStation D1000 Screen Tape (#5067-5582,5583, Agilent Technologies). We finally clonally amplified the quantified libraries on a cBOT and sequenced them on the HiSeq X with 150bp paired end chemistry.

### Whole Genome Re-sequencing and variant discovery

We trimmed the raw reads using TRIMMOMATIC^87^ to have an mean PHRED-scaled quality of 30 in a sliding window of 15 bp, and removed any read that was shorter than 36 bp after trimming from further analysis. We aligned these reads to a Bengal tiger reference genome (JAHFZI000000000) using BOWTIE2^88^. The alignments were then saved in a binary format (BAM) using SAMTOOLS1.9^89^. We marked duplicate reads with the Picard Tools ‘MarkDuplicates’ command (http://broadinstitute.github.io/picard). We called variants from the BAM files using Strelka with default options^90^. The variants were filtered with VCFtools^91^ to retain biallelic sites with a minimum minor allele count of 3, remove indels and loci with mean depth across individuals below 2.5 percentile and above 97.5 percentile across all loci. We removed all sites with missingness > 20% after removing genotypes with genotype quality (GQ) less than 30. We removed the X chromosome from the analysis. 1601148 SNPs with mean read depth = 17.2 (+/-1.8) remained for analyses after these filtering steps.

### Runs of Homozygosity

We identified ROH using a sliding window, likelihood ratio method^92,93^. For this analysis we used SNP data without the minimum genotype quality filter described above, and then used genotype likelihoods, rather than called genotypes, as input in order to maximize the genomic information available for detecting ROH. We split each chromosome into sliding windows of 200 SNPs and step size of 10 SNPs. For each individual *i* and window *j*, we calculated the probability of the observed genotype likelihood at each SNP *k* (*G*_*k*_) assuming the two alleles were IBD, and also assuming they were non-IBD (Wang et al. 2009). We calculated the LOD score for each window as

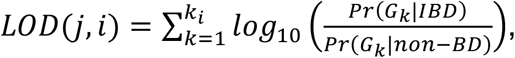

LOD scores across all windows and individuals form a bimodal distribution. Windows with LOD score ≥ 0 were called IBD, while windows in the left hand side of the distribution were called non-IBD as in Kardos et al.^44,94^. Consecutive IBD windows were joined to map the starting and stop positions of ROH for each individual. *F*_ROH_ was calculated as the fraction of the autosomal genome in ROH for example,

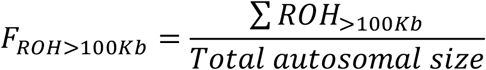

We calculated the percent of the genome in ROH above a particular size class to estimate inbreeding arising from ancestors in different historical time periods. The coalescent times of ROH were estimated as *g=100/(2rL)* where *g* is the expected time (in generations) back to the parental common ancestor where the IBD haplotypes forming an ROH coalesce^95^, *r* is the recombination rate (here, 1.9cM per Mb based on domestic cat^45^) and *L* is the length of the ROH in Mb. For example, a 100Kb ROH are estimated to arise from a single haplotype in an ancestor 263 generations ago on average. An individual’s genomic inbreeding arising from ancestors up to 263 generations ago is then estimated using only ROH longer than 100 kb. Similar calculations were performed for other ROH lengths. Table 1 lists the lengths of ROH and corresponding average estimated coalescent times for the IBD haplotypes that form the ROH.

**Table 1.** ROH lengths and the corresponding expected mean time of their origin due to inbreeding where number of years assumes a generation length of 5 years.

| ROH length(s) | generations since parents shared ancestor | years since parents shared ancestor |
| --- | --- | --- |
| <100Kb | More than 263 | More than 1315 |
| >100Kb | Up to 263 | Up to 1315 |
| >1Mb | Up to 26 | Up to 130 |
| >5Mb | Up to 5 | Up to 25 |
| >10Mb | Up to 3 | Up to 15 |

We constructed percentile bootstrap confidence intervals for mean *F*_ROH_ in each population^96^. We randomly resampled individuals within a population with replacement 10,000 times, each time calculating the mean *F*_ROH_ as described above. The 0.025 and 0.975 quantiles defined the 95% confidence interval for mean *F*_ROH_ in each population. We used a randomization procedure to test for a difference in the mean *F*_ROH_ for each pair of populations as follows. First, we randomized the population affiliation for each individual 10,000 times. For each randomization replicate, we calculated the mean *F*_ROH_ for each of the two populations. We calculated the *P*-value as the proportion of 10,000 randomization replicates where the difference in *F*_ROH_ was at least as large as the empirical difference in *F*_ROH_.

### IBD stretches of genome shared between pairs of individuals

We estimated stretches of genome identical by descent (IBD) between pairs of individuals using *IBDseq* default parameters^97^. We filtered for shared stretches longer than 1Mb and estimated sum of shared lengths of more than 1Mb per autosomal genome size. We estimated this for all pairs of individuals within a population and all pairs across populations.

### Estimating effective population size from ROH

We used the estimates of *F*_ROH_ using ROH arising from ancestors in 4 different estimated time frames (Figure 1) to estimate *N*_e_ over each respective time frame. If the estimated coalescent times for ROH are unbiased, then the average *F*_ROH_ based on ROH with estimated maximum coalescent times less than *t* generations back in time is an estimator of the inbreeding accumulated in the population from the time of sampling back to *t* generations ago. We estimated *N*_e_ from the following expression of mean expected individual inbreeding as a function of *N*_e_ over *t* generations

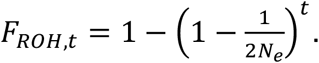

This approach assumes that the probability of inferring an ROH segment due to several smaller homozygous segments is very low. We calculated 95% confidence intervals for *N*_e_ by bootstrapping over individuals within each population as described above for our estimates of mean *F*_ROH_.

### Mitochondrial haplotype diversity

We used the reads aligned to the mitochondrial scaffold of BenTig1.0. We called consensus fasta files for each individual from this as described in Khan et al.^81^. These multiple consensus fasta files were aligned using clustal omega (https://www.ebi.ac.uk/Tools/msa/clustalo/) and written to nexus output files. We then used DnaSP 6^98^ to estimate haplotype diversity.

### Identifying effect of mutations

We re-filtered the SNPs as described above with VCFtools^91^, except here we removed genotypes with phred-scaled genotype quality <30, and then removed loci that were monomorphic, had >20% missing data, or read depth in the bottom and top 2.5% quantiles for mean read depth across individuals. We also removed loci and with extreme values of *F*_IS_ (*F*_IS_<-0.5 or *F*_IS_ >0.95) estimated across all Indian tigers to remove loci likely frequent genotyping or read mapping errors; this expectedly resulted in mean *F*_IS_>0 (0.14).

We used Ensembl Variant Effect Predictor^59^ on SNPs to identify consequence of a mutation and classify them as missense, loss-of-function (LOF) or intergenic as described by Xue et al.^28^. We used annotation files generated for the Bengal tiger assembly using RNA seq.

### Mutation load

We used the R_XY_ method described by Do et al.^62^ and implemented by Xue et al.^28^ to estimate relative excess of mutation loads in a population X with respect to another population Y. We randomly selected 7 individuals from each population and estimated the relative mutation load for each of the three possible pairwise combinations of the S-I, s-L-C, c-L-C populations. Standard deviations were obtained by 100 rounds of jackknife. For each round of jackknife for missense and LOF mutations, we randomly excluded 10-15% of loci and repeated the estimation. For intergenic variants, 100,000 loci were randomly excluded for each round.

To infer the ancestral state of each site in the tiger reference we converted the references for the Domestic cat (GenBank: GCA_000181335.4), Cheetah (GCA_003709585.1) and Lion (GCA_008795835.1) into FASTQ reads by sliding across the genome in non-overlapping windows of 100 base pairs and transforming each window into a separate FASTQ read. The resulting FASTQ reads were then mapped to the tiger reference genome with bwa mem v0.7.17, slightly lowering the mismatch penalty (*-B* 3) and removing reads that mapped to multiple regions. Mapped reads were realigned around indels using GATK *IndelRealigner*. Next, we converted the mapped reads into a haploid FASTA consensus sequence, excluding all sites with depth above one (as such sites contain at least one mismapped read) using ANGSD *-dofasta*. The ancestral allele at a locus was then determined as the majority allele found in genomic alignment of the domestic cat, cheetah and lion. Sites where the ancestral allele could not be identified (e.g., where the ancestral identified in the cat-cheetah-lion alignment was not present in our sample of tigers) were excluded.

We counted the number of homozygous deleterious alleles per individual as number of derived missense and LOF mutations. We quantified uncertainty in the mean number of homozygous deleterious alleles for each population using percentile bootstrap confidence intervals as described above. We used randomization tests as described above to test for differences in the mean number of homozygous damaging and LOF alleles per individual for each pair of populations.

### Site-frequency spectrum

We equalized the sample sizes across the populations, and across loci within each population by randomly subsampling 16 non-missing alleles from each locus in each population population before estimating the derived neutral and derived damaging allele frequencies. For the SFS analysis of all populations combined, we subsampled 88 non-missing alleles from each locus. Repeating the subsampling process and analysis had no effect on the results.

We used randomization tests to determine if the mean putatively damaging derived allele frequency was statistically significantly different from the mean putatively neutral derived allele frequency in each population, and for all populations combined (Figure 3). We randomly reassigned the neutral versus damaging status for each allele frequency 5,000 times. For each of the 5,000 replicates, we recalculated the randomized mean derived neutral and damaging allele frequency. A *P*-value was calculated as the proportion of the 5,000 permutations that resulted in a difference in the mean neutral versus damaging derived allele frequency that was larger than observed in the non-randomized data.

### Genetic rescue and F_ST_

We estimated genome wide genetic differentiation as *F*_ST_ with VCFtools. For loci with high frequency (>0.85, closer to fixation) deleterious (missense and LOF) alleles in a population, we listed all the loci in each target population (population in need of rescue). For every possible source population, we estimated the proportion of these loci with frequency < 0.85. We then calculated *F*_ST_ at only these loci to estimate damaging loci *F*_ST_. The proportion or loci rescued was calculated as the fraction of loci that had < 0.85 frequency in the possible source population.

We investigate this for five possible source populations, including tigers from Kanha Tiger Reserve (KTR), Wayanad Wildlife Sanctuary (WAY), Ranthambore Tiger Reserve (RTR), Corbett Tiger Reserve (COR) and Kaziranga Tiger Reserve (KAZ).

## Acknowledgements

This work was supported primarily by a DBT/Wellcome Trust India Alliance Fellowship [IA/S/16/2/502714] awarded to UR. AK received support from SciGenome Research Foundation. Permissions for sample collection were granted in letters F-19 ()CWLW/2018-19/653-55 dated 15-06-2018; E2/PCCF/WL/CR/40/2013-14 and PCCF(WL)/E2/CR/28/2017-18. Zoo samples were collected as described in Sagar et al., 2021, under CZA permit - 9-3/2005-CZA(Vol lll)(D)/694/2017. Leslie Lyons collected and kindly shared the list of cat mutations. Gratitude to NCBS IT team support during data analysis; NCBS data cluster used is supported under project no. 12-R&D-TFR-5.04-0900, Department of Atomic Energy, Government of India). Byron Weckworth, Meghana Natesh, Mousumi Ghosh, Megan Aylward, Chris Kyriazis, Ullas Karanth and Phil Hedrick provided helpful discussions and comments on drafts of the manuscript. Gratitude to field assistants for their continuous presence in the field and assisting in sample collection for this work. We thank the editor and two anonymous reviewers for their comments that helped improve the manuscript.

## Data availability

All sequencing data have been deposited in BioProject PRJNA728665 in NCBI

## Notes

### Competing Interest Statement

The authors have declared no competing interest.

### Summary of Updates

This update includes additional analysis on deleterious alleles, SFS and genetic rescue

