## Supplementary material for "Genomic evidence for inbreeding depression and purging of deleterious genetic variation in Indian tigers"

**Supplementary Methods:**

*Mapping to domestic genome and variant discovery for predicting potential diseases*

We mapped trimmed FASTQ reads to the *Felis catus* reference genome version 9 (GenBank: GCA_000181335.4) using BWA-MEM (Li 2013). The alignments were then saved in a binary format (BAM) using SAMTOOLS1.9 (Li et al., 2009). We marked duplicate reads with the Picard Tools `MarkDuplicates` command (http://broadinstitute.github.io/picard). We called variants from the BAM files using Strelka with default options (Saunders 2012). The variants were filtered with VCFtools (Danecek et al. 2011) to retain biallelic sites with a minimum minor allele count of 3, minimum base quality 30, minimum depth 10, minimum genotype quality 30, removed Indels and had less than 20% missing data.

*Identifying effect of mutations*

We used Ensembl Variant Effect Predictor (McLaren et al. 2016) on SNPs identified with domestic cat reference genome to identify consequence of a mutation and classify them as missense, loss-of-function (LOF) or intergenic as described by Xue et al. (2015). We then compared these mutations with an annotated dataset of known disease cause mutations in domestic cats.

*GERP based Mutation load*

We estimated GERP scores for each locus on our reference tiger genome as described in van der Valk et al. (2019). We selected the loci with the top 0.1 percent GERP scores from the variant call file filtered from the tiger genome. These are potentially loci hosting highly deleterious alleles. We selected the loci with the least 1 percent GERP scores from the same file. These are potentially neutral loci. Next we used the R_XY_ method described by Do et al. (2015) and implemented by Xue et al. (2015) to estimate relative excess of mutation loads as described in the main text.

**Supplementary table 1:** Sample sequencing and location. north-west Indian: 1=Ranthambore Tiger Reserve (n=11), 2=Sariska Tiger Reserve (n=7), south Indian population: 3=Bandipur Tiger Reserve (n=2), 4=Wayanad Wildlife Sanctuary (n=6), 5=Periyar Tiger Reserve (n=1), central Indian Population: 6=Kanha Tiger Reserve (n=8), 7=Corbett Tiger Reserve (n=2), 8=Lalgarh Range (n=1), 10=Sunderban Tiger Reserve (n=2), 11=Bor Tiger Reserve (n=1),north-east Indian population: 9=Kaziranga Tiger Reserve (n=3) , Zoo: Zoo=Nanadankanan Zoo (n=5).

| Individual | Location | Sequencing depth | F_PED_ | Proportion/genome [>100Kb] | Proportion/genome [>1Mb] | Biosample accession |
| --- | --- | --- | --- | --- | --- | --- |
| BEN_CI15 | 6 | 24 |  | 0.316 | 0.091 | [SAMN20398718](https://dataview.ncbi.nlm.nih.gov/object/SAMN20398718)^1^ |
| BEN_CI16 | 6 | 15 |  | 0.476 | 0.304 | [SAMN20398719](https://dataview.ncbi.nlm.nih.gov/object/SAMN20398719)^1^ |
| BEN_CI18 | 6 | 26 |  | 0.303 | 0.080 | [SAMN20398720](https://dataview.ncbi.nlm.nih.gov/object/SAMN20398720)^1^ |
| BEN_CI19 | 6 | 13 |  | 0.346 | 0.146 | [SAMN20398721](https://dataview.ncbi.nlm.nih.gov/object/SAMN20398721)^1^ |
| BEN_CI2 | 14 | 8 |  | 0.398 | 0.207 | SAMN17487589^2^ |
| BEN_CI21 | 11 | 18 |  | 0.411 | 0.197 | [SAMN20398722](https://dataview.ncbi.nlm.nih.gov/object/SAMN20398722)^1^ |
| BEN_CI3 | 6 | 22 |  | 0.345 | 0.123 | SAMN17487590^2^ |
| BEN_CI4 | 6 | 13 |  | 0.332 | 0.125 | SAMN17487591^2^ |
| BEN_CI5 | 6 | 15 |  | 0.293 | 0.063 | SAMN17487592^2^ |
| BEN_CI6 | 6 | 11 |  | 0.328 | 0.105 | SAMN17487593^2^ |
| BEN_CI7 | 6 | 8 |  | 0.428 | 0.253 | SAMN17487594^2^ |
| BEN_NE1 | 9 | 22 |  | 0.330 | 0.097 | SAMN17487596^2^ |
| BEN_NE2 | 9 | 18 |  | 0.605 | 0.469 | SAMN17487597^2^ |
| BEN_NE3 | 9 | 12 |  | 0.523 | 0.363 | SAMN17487598^2^ |
| BEN_NE4 | 9 | 8 |  | 0.373 | 0.229 | [SAMN20398723](https://dataview.ncbi.nlm.nih.gov/object/SAMN20398723)^1^ |
| BEN_NOR_SJ1 | 13 | 12 |  | 0.414 | 0.202 | SAMN09080471^3^ |
| BEN_NOR1 | 7 | 23 |  | 0.400 | 0.159 | SAMN17487599^2^ |
| BEN_NOR2 | 7 | 24 |  | 0.453 | 0.234 | SAMN17487600^2^ |
| BEN_NW10 | 1 | 27 |  | 0.620 | 0.519 | [SAMN20398724](https://dataview.ncbi.nlm.nih.gov/object/SAMN20398724)^1^ |
| BEN_NW11 | 2 | 23 |  | 0.495 | 0.360 | [SAMN20398725](https://dataview.ncbi.nlm.nih.gov/object/SAMN20398725)^1^ |
| BEN_NW12 | 2 | 16 |  | 0.608 | 0.499 | [SAMN20398726](https://dataview.ncbi.nlm.nih.gov/object/SAMN20398726)^1^ |
| BEN_NW13 | 2 | 23 |  | 0.577 | 0.470 | [SAMN20398727](https://dataview.ncbi.nlm.nih.gov/object/SAMN20398727)^1^ |
| BEN_NW14 | 2 | 23 |  | 0.586 | 0.478 | [SAMN20398728](https://dataview.ncbi.nlm.nih.gov/object/SAMN20398728)^1^ |
| BEN_NW17 | 1 | 14 |  | 0.556 | 0.430 | [SAMN20398729](https://dataview.ncbi.nlm.nih.gov/object/SAMN20398729)^1^ |
| BEN_NW18 | 1 | 21 |  | 0.633 | 0.530 | [SAMN20398730](https://dataview.ncbi.nlm.nih.gov/object/SAMN20398730)^1^ |
| BEN_NW19 | 1 | 23 |  | 0.524 | 0.386 | [SAMN20398731](https://dataview.ncbi.nlm.nih.gov/object/SAMN20398731)^1^ |
| BEN_NW20 | 1 | 18 |  | 0.554 | 0.445 | SAMN12620966^4^ |
| BEN_NW24 | 1 | 23 |  | 0.569 | 0.454 | [SAMN20398733](https://dataview.ncbi.nlm.nih.gov/object/SAMN20398733)^1^ |
| BEN_NW3 | 2 | 20 |  | 0.609 | 0.495 | SAMN17487602^2^ |
| BEN_NW4 | 2 | 20 |  | 0.592 | 0.490 | SAMN17487603^2^ |
| BEN_NW5 | 1 | 34 |  | 0.483 | 0.342 | SAMN12549969^4^ |
| BEN_NW55 | 2 | 18 |  | 0.571 | 0.456 | [SAMN20398735](https://dataview.ncbi.nlm.nih.gov/object/SAMN20398735)^1^ |
| BEN_NW57 | 1 | 15 |  | 0.625 | 0.519 | [SAMN20398736](https://dataview.ncbi.nlm.nih.gov/object/SAMN20398736)^1^ |
| BEN_NW6 | 1 | 15 |  | 0.491 | 0.356 | [SAMN20398737](https://dataview.ncbi.nlm.nih.gov/object/SAMN20398737)^1^ |
| BEN_NW63 | 1 | 15 |  | 0.632 | 0.529 | [SAMN20398738](https://dataview.ncbi.nlm.nih.gov/object/SAMN20398738)^1^ |
| BEN_NW8 | 1 | 16 |  | 0.589 | 0.474 | [SAMN20398739](https://dataview.ncbi.nlm.nih.gov/object/SAMN20398739)^1^ |
| BEN_NW9 | 1 | 21 |  | 0.593 | 0.482 | [SAMN20398740](https://dataview.ncbi.nlm.nih.gov/object/SAMN20398740)^1^ |
| BEN_SI1 | 4 | 21 |  | 0.461 | 0.249 | SAMN17487604^2^ |
| BEN_SI11 | 4 | 12 |  | 0.423 | 0.220 | [SAMN20398741](https://dataview.ncbi.nlm.nih.gov/object/SAMN20398741)^1^ |
| BEN_SI18 | 3 | 16 |  | 0.436 | 0.232 | [SAMN20398742](https://dataview.ncbi.nlm.nih.gov/object/SAMN20398742)^1^ |
| BEN_SI19 | 3 | 17 |  | 0.447 | 0.238 | [SAMN20398743](https://dataview.ncbi.nlm.nih.gov/object/SAMN20398743)^1^ |
| BEN_SI2 | 3 | 6 |  | 0.432 | 0.244 | SAMN17487605^2^ |
| BEN_SI3 | 4 | 16 |  | 0.487 | 0.287 | SAMN17487606^2^ |
| BEN_SI5 | 5 | 10 |  | 0.668 | 0.532 | SAMN17487608^2^ |
| BEN_SI6 | 4 | 20 |  | 0.479 | 0.280 | SAMN17487609^2^ |
| BEN_SI8 | 4 | 25 |  | 0.552 | 0.376 | SAMN17487607^2^ |
| BEN_SI9 | 4 | 23 |  | 0.442 | 0.218 | [SAMN20398744](https://dataview.ncbi.nlm.nih.gov/object/SAMN20398744)^1^ |
| BEN_SI_SJ1 | 12 | 11 |  | 0.423 | 0.226 | SAMN09080449^3^ |
| BEN_SI_SJ2 | 12 | 11 |  | 0.370 | 0.144 | SAMN09080448^3^ |
| BEN_SU2 | 10 | 15 |  | 0.470 | 0.273 | [SAMN20398745](https://dataview.ncbi.nlm.nih.gov/object/SAMN20398745)^1^ |
| BEN_SU3 | 10 | 41 |  | 0.565 | 0.403 | [SAMN20398746](https://dataview.ncbi.nlm.nih.gov/object/SAMN20398746)^1^ |
| LGS1 | 8 | 15 |  | 0.408 | 0.208 | [SAMN20398747](https://dataview.ncbi.nlm.nih.gov/object/SAMN20398747)^1^ |
| ZSB_01 | Zoo | 20 | 0.21 | 0.389 | 0.190 | SAMN20424160^5^ |
| ZSB_02 | Zoo | 20 | 0.28 | 0.482 | 0.329 | SAMN20424161^5^ |
| ZSB_03 | Zoo | 29 | 0.28 | 0.468 | 0.307 | SAMN20424162^5^ |
| ZSB_04 | Zoo | 26 | 0.28 | 0.466 | 0.309 | SAMN20424163^5^ |
| ZSB_05 | Zoo | 30 | 0.26 | 0.451 | 0.281 | SAMN20424164^5^ |

1) This study
2) E. E. Armstrong, *et al.*, Recent Evolutionary History of Tigers Highlights Contrasting Roles of Genetic Drift and Selection. *Mol. Biol. Evol.* **38**, 2366–2379 (2021).
3) Y.-C. Liu, *et al.*, Genome-Wide Evolutionary Analysis of Natural History and Adaptation in the World’s Tigers. *Current Biology* **28**, 3840–3849.e6 (2018).
4) A. Khan, *et al.*, Are shed hair genomes the most effective noninvasive resource for estimating relationships in the wild? *Ecol. Evol.* **10**, 4583–4594 (2020).
5) V. Sagar, *et al.*, High frequency of an otherwise rare phenotype in a small and isolated tiger population. *PNAS* (in press manuscript ID 2020-25273).

**Supplementary Table 2:** Genes with derived missense mutations and potential diseases predicted with domestic cat annotations

| **Gene** | **Disease** | **Reference** |
| --- | --- | --- |
| MYBPC3 | Hypertrophic Cardiomyopathy | Meurs et al. 2005, Meurs et al. 2007 |
| MYBPC2 | Hypertrophic Cardiomyopathy | Meurs et al. 2005, Meurs et al. 2007 |
| WNK4 | Hypokalemia | Gandolfi et al. 2012 |
| CEP290 | Progressive Retinal Atropy | Menotti-Raymond et al. 2010 |
| PKD1 | Polycystic Kidney Disease | Lyons et al. 2004 |
| DPYS | Dihydropyrimidiname Deficiency | Chang et al. 2012 |
| SLC3A1 | Cystinuria, Type 1A | Mizukami et al. 2015 |
| HEXB | Gangliosidosis | Martin et al. 2004, Kanae et al. 2007 |
| GRHPR | Hyperoxaluria | Goldstein et al. 2009 |
| TPO | Hypothyroidism | Giger et al. 2015 |
| GNPTAB | Mucolipidosis II | Mazrier et al. 2003 |
| GUSB | Mucopolysaccharidosis VII | Fyfe et al. 1999, Wang et al. 2015 |
| UROS | Porphyria | Clavero et al. 2010, Clavero et al. 2013 |

**a**

**b**

Number of mitochondrial haplotypes

|  | number of haplotypes |
| --- | --- |
| small isolated | 4 |
| southern large connected | 5 |
| central large connected | 7 |
| All India | 20 |

Mitochondrial haplotype diveristy

|  | Haplotype diversity |
| --- | --- |
| small isolated | 0.455 |
| southern large connected | 0.833 |
| central large connected | 1 |
| All India | 0.909 |

**Supplementary figure 1:** (a) Number of mitochondrial haplotypes detected and (b) mitochondrial haplotype diversity

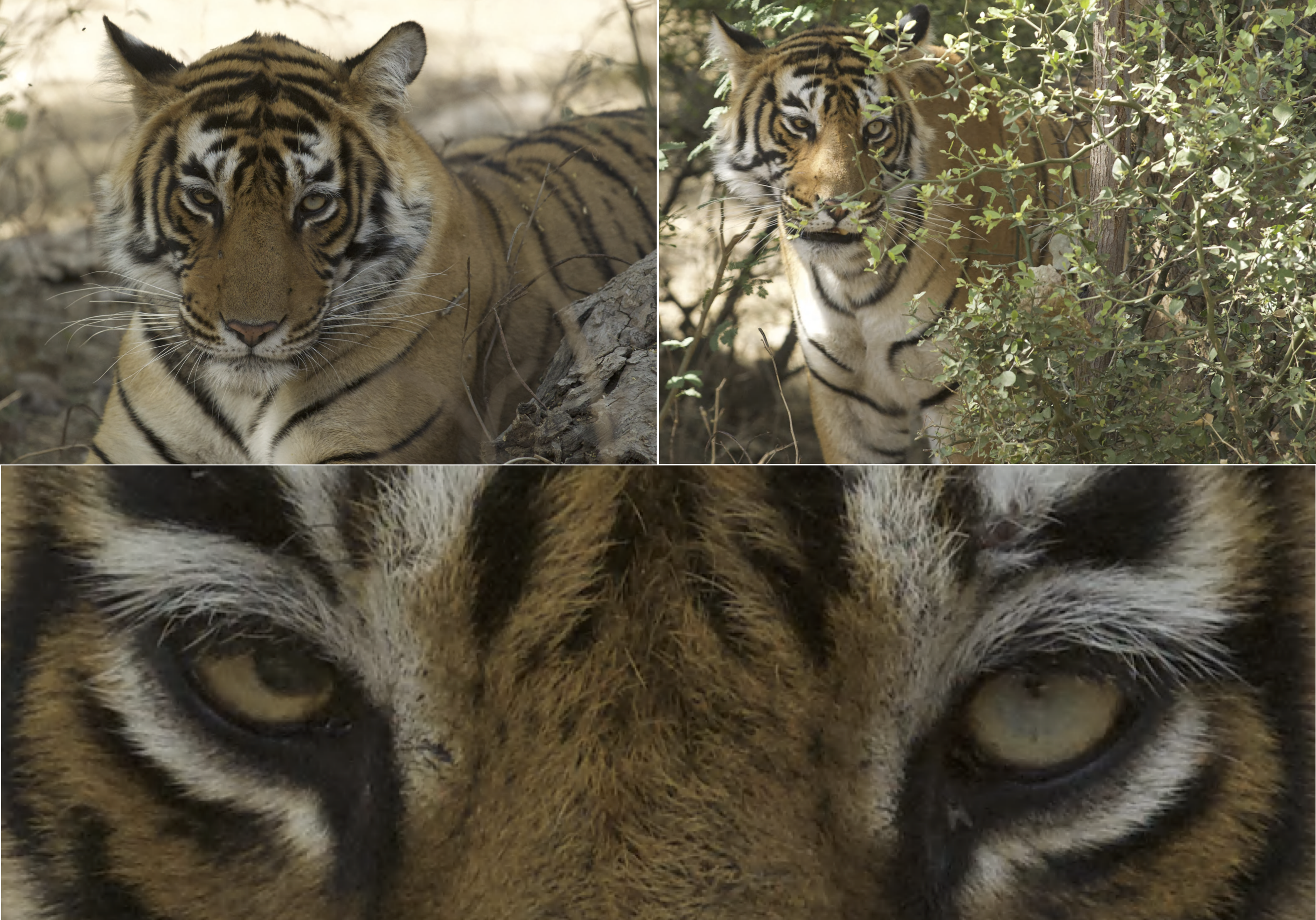

**c**

**b**

**a**

**Supplementary figure 2:** Photos of tigeress T99 from Ranthambore Tiger Reserve (a,b). Close up of her eyes (c). Her litter-mate BEN_NW63 has 35% of her genome in more than 10Mb long ROH.

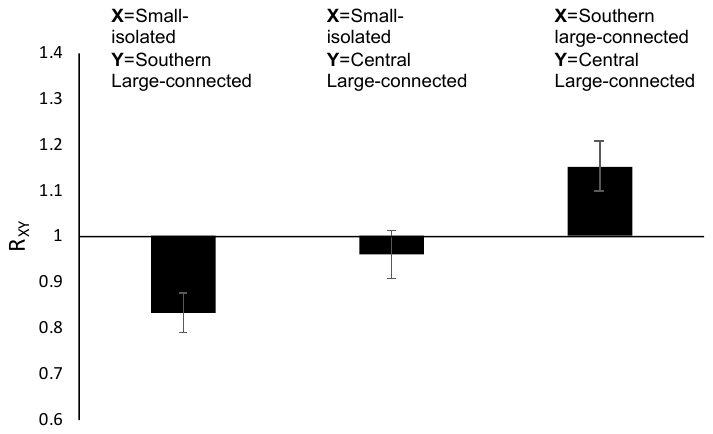

**Supplementary figure 3:** Relative mutation loads based on loci with 0.1 percent of the top GERP scores. Error bars are standard errors from jacknife.
